# Loss of REP1 impacts choroidal melanogenesis in choroideremia

**DOI:** 10.1101/2023.07.20.549875

**Authors:** Hajrah Sakar, Dhani Tracey-White, Ahmed M. Hagag, Thomas Burgoyne, Lasse D. Jensen, Malia M. Edwards, Mariya Moosajee

## Abstract

Choroideremia (CHM) is a rare X-linked chorioretinal dystrophy affecting the photoreceptors, retinal pigment epithelium (RPE) and choroid, however, the involvement of the choroid in disease progression is not fully understood. CHM is caused by mutations in the *CHM* gene, encoding the ubiquitously expressed Rab escort protein 1 (REP1). REP1 plays an important role in intracellular trafficking of vesicles, including melanosomes. In this study, we examined ultrastructure of the choroid in *chm^ru848^* fish and *Chm^null/WT^* mouse models using transmission electron microscopy. Significant pigmentary disruptions were observed, with a lack of melanosomes in the choroid of *chm^ru848^*fish from 4 days post fertilisation (4dpf). Total melanin and expression of melanogenesis genes *tyr*, *tryp1a*, *mitf, dct* and *pmel* were also reduced from 4dpf. In *Chm^null/WT^* mice, choroidal melanosomes were significantly smaller at 1 month and at 1 year, eumelanin was reduced, and the choroid was thicker. The choroid in CHM patients was also examined using optical coherence tomography (OCT) and OCT-angiography (OCT-A) and the area of preserved choriocapillaris (CC) was found to be smaller than that of overlying photoreceptors, indicating that the choroid is degenerating at a faster rate. Histopathology of an enucleated eye from a 74-year-old CHM male patient revealed isolated areas of RPE but no associated underlying CC. Significant degenerative changes in the choroid of CHM patients and animal models are seen, highlighting the importance of administrative routes involving the choroid, such as suprachoroidal delivery. Pigmentary disruptions in CHM animal models reveal an important role for REP1 in melanogenesis, and drugs that improve melanin production represent a potential novel therapeutic avenue.

## Introduction

Choroideremia (CHM) is a rare X-linked inherited chorioretinal dystrophy, caused by mutations in the *CHM* gene, which encodes the ubiquitously expressed Rab escort protein 1 (REP1). Affected males typically present with nyctalopia in early childhood, progressing to constriction of visual fields and eventual loss of central vision by middle-late age. CHM is characterised by progressive degeneration of the photoreceptors, retinal pigment epithelium (RPE) and choroid; however, the involvement of the choroid is not yet fully understood.

REP1 is responsible for the prenylation of Rab GTPases and plays an important role in intracellular trafficking of vesicles. Rab27a, a known target of REP1, is essential for melanosome movement. Rab27a, which binds to melanosomes, forms a tripartite complex with myosin VIIa, on actin filaments, and the linker protein MyRIP [1–3], allowing movement of the melanosomes along the actin cytoskeleton. The *Chm^Flox^, Tyr-Cre+* mouse model has previously been shown to have reduced melanosomes in the apical processes of the RPE, however no differences were detected in melanosome distribution in the choroid [4]. Melanosomes are the site of melanin synthesis and storage; and protect the retina from photo-oxidative stress by absorbing light and scavenging reactive oxygen species. There are two types of melanin in mammals, eumelanin which is the black/brown pigment and acts as an antioxidant, and pheomelanin which is the red/yellow pigment and is pro-oxidant [5]. RPE is mostly eumelanin whereas the choroid contains both eumelanin and pheomelanin, with the pheomelanin content varying depending on eye colour [6].

In this paper, we investigate the involvement of the choroid in the progression of CHM. The ultrastructure of the choroid, with particular focus on the melanosomes, is examined in two CHM animal models. Firstly, the *chm^ru848^* zebrafish, which due to lack of a REP2 orthologue (which in humans compensates for REP1 loss in most tissues except the retina), displays systemic degeneration with a mean survival of 4.8 days post fertilisation (dpf). *chm^ru848^* zebrafish have small eyes with loss of iridophores and cataracts, the retina shows early signs of patchy photoreceptor cell loss, followed by RPE atrophy and areas of hypertrophy with invasion into the inner nuclear layer (INL), rapidly progressing to widespread cell death and loss of retinal lamination [7]. Secondly, the *Chm^null/WT^* mouse model, a heterozygous female carrier model with a single *Chm^null^* allele, which displays a progressive retinal degeneration with areas of hypopigmentation throughout the whole retina from 1 month, which become confluent by 4 months. Progressive thinning of the outer nuclear layer (ONL) and patchy depigmentation of the RPE is observed from 2 months, which expand to severe degeneration by 8 months [8]. We then examine the choroid in CHM patients; the relationship between the CC and photoreceptors in CHM patients is studied using optical coherence tomography (OCT) and OCT-angiography (OCT-A), with comparison to the ultrastructural analysis of an enucleated eye from a 74 year old Caucasian male CHM patient [9]. Altogether, this provides an extensive overview into the pathophysiological role of the choroid in CHM.

## Methods

### Zebrafish strains and husbandry

Wild-type AB (wt) and choroideremia (*chm^ru848^*) zebrafish were generated by natural pair-wise matings of genotyped heterozygous fish and raised at 28.5°C on a 14 h light/10 h dark cycle under the Animals Scientific Procedures Act at the UCL Bloomsbury campus zebrafish facility. The zebrafish reporter line *tg(fli1a:egfp)* was sourced from EZRC and crossed with the *chm^ru848^* line to create a tagged line.

### Mouse strains and husbandry

All mouse samples used in this study were kindly donated by Professor Miguel Seabra, Imperial College London. The schedule 1 procedure and collection of retinas was completed adhering to the ARVO Statement for the Use of Animals in Ophthalmic and Vision Research. The conditional knock-out mouse line *Chm^null/WT^* was generated previously and genotyping of mice was performed as described [8]. As controls, female *Chm^flox/flox^* littermates were used.

### Zebrafish retinal microangiography

Zebrafish larvae were euthanized in 0.04% MS-222 (Ethyl 3-aminobenzoate methane sulfonic acid salt 98%, Sigma Aldrich) for 15 min and fixed for 30 min at room temperature in 4% paraformaldehyde (PFA). Retinal flat mounts were prepared and the choriocapillaris was visualised and analysed as previously described [10]. Briefly, the larvae were enucleated under a dissection microscope (Nikon SMZ 1500) and retinae were prepared as flat mounts with the choroidal side up in Vectashield mounting medium (H-1000 Vector laboratories) and imaged by confocal microscope (LSM 700 Zeiss Upright confocal). Image analysis was done using Photoshop CS6 (Adobe) or ImageJ (NIH).

### Zebrafish and mouse histology and transmission electron microscopy (TEM)

Histology analysis of mouse retinas was achieved by taking semi-thin sections (∼500um thick) during the TEM processing protocol and after staining with 0.05% toluidine blue, sections were imaged using a light microscope. For TEM, samples were fixed in 2% paraformaldehyde-2% glutaraldehyde prior to incubation with 2% osmium tetroxide-1.5% potassium ferrocyanide. Following dehydration in an ethanol series and propylene oxide, samples were embedded in EPON 812 resin. Using a Leica EM UC7 ultramicrotome, ultrathin 70nm sections were cut, collected on formvar-coated copper slot grids (EMS) and stained with lead citrate (Agar Scientific). TEM Sections were examined on a JEOL 1400+ fitted with AMT side mount camera. A macro was made for ImageJ to measure the area of individual melanosomes from images taken at 6,000x magnification. Initially, the macro enhances image contrast before making a binary image. Subsequently, functions are run in ImageJ to fill in missing pixel within melanosome structures. A filter is applied to reduce background and the ‘Analyze Particles’ feature used to measure the area of any structures greater than 400 pixel^2^ (to exclude background) and that have a circularity of 0.2 – 1.0 (to exclude most of the irregular shaped structure caused by overlapping melanosomes). Value above 0.5 µm^2^ were removed from the analysis, to avoid including any potential overlapping melanosomes that were measured.

### Zebrafish whole melanin quantification

Melanin content was determined according to the protocol by Agalou et al. [11]. Briefly, eyes from 20 fish were sonicated in cold lysis buffer (20 mM sodium phosphate (pH 6.8), 1% Triton X-100, 1 mM EDTA, 1x Halt protease and phosphatase inhibitors cocktail). An aliquot of the lysate was reserved to determine protein content using Pierce BCA protein kit. The lysate was centrifuged at 10,000 × *g* for 10 min. The pellet was resuspended in 1 mL 1 N NaOH/10% DMSO and incubated at 95°C for 1 hour. Absorbance was measured at 405 nm. Data were normalized to total protein content.

### HPLC analysis of eumelanin and pheomelanin in mouse samples

RPE and choroid were dissected and snap frozen until analysis. Melanin was analysed according to the protocol by Affenzeller et al., [12] with slight modifications. Samples were treated with 10 µL proteinase K (10 mg/mL) in 500 µL TRIS-HCl buffer (1 M, pH 8.0) for 2 hours at 55°C in a shaker. Treatment was stopped by the addition of 300 µL 6M HCl and centrifuged at 13,000 rpm for 15 min. Samples were then oxidised with 100 μL H_2_O, 375 μL K_2_CO_3_ (1 M) and 25 μL H_2_O_2_ (30%) for 20 h at 25°C with vigorous shaking. After this time any remaining H_2_O_2_ was inactivated by the addition of 50 µL Na_2_SO_3_ (10% (w/v) and 140 µL HCl (6 M). Samples were then centrifuged at 13,000 rpm for 30 min and the supernatant was transferred into a fresh tube. Oxidised samples were treated by solid phase extraction on Strata^™^-X 33 μm Polymeric Reversed Phase cartridges 30 mg/3 mL (Phenomenex, Torrance, USA). HPLC analysis was performed using a Waters ACQUITY® UPLC® equipped with a Waters ACQUITY 2996 PDA detector. The stationary phase was Waters ACQUITY UPLC BEH C18 1.7 μm, 2.1 x 50 mm column. The analysis was 32 minutes, with a flow rate of 0.062 ml/min and a 10 μl injection. The elution method used was a gradient of water, 0.1% formic acid and acetonitrile; 0-16 min 1% acetonitrile, 16-24 min up to 95% acetonitrile, 24-28 min 95% acetonitrile, 28-28.8 min 95 to 1% acetonitrile, 28.8-32 min 1% acetonitrile. Mass spectra was acquired in negative ion mode. Eumelanin was quantified by detection of the oxidation products PDCA and PTCA at 154 and 198 m/z. Pheomelanin was quantified by detection of oxidation products TDCA and TTCA at 128 and 172 m/z.

### RT-qPCR

Total RNA was extracted from samples using the RNeasy mini kit (QIAGEN) from dissected eyes of 10 fish. cDNA was synthesised from 1 μg of RNA using the Superscript II First Strand cDNA synthesis kit (Invitrogen). Transcript levels were analysed using SYBR Green MasterMix (ThermoFisher) on a StepOne Real-Time PCR system (Applied Biosystems), under standard cycling conditions and normalised to housekeeping gene B-actin. All samples were assayed in triplicate. Primer sequences are shown in Table S1.

### OCT/OCT-angiography

Analysis of a natural history study was performed to investigate the relationship between choriocapillaris (CC) and photoreceptors in patients with molecularly confirmed choroideremia. A commercially available spectral-domain OCT machine (RTVue-XR, OptoVue) was used to acquire simultaneous 6x6 mm macular structural OCT and OCT angiography scans as part of a prospective longitudinal study at Moorfields Eye Hospital (London, UK). Volumetric OCT and OCT-A were segmented and processed to construct *en face* images of the EZ and CC layers, respectively. Areas of preserved EZ and CC were manually delineated and calculated. Detailed description of image processing and area measurement methods, as well as reliability analyses were reported in our previous publication [13]. The absolute and percentage differences between CC and EZ preserved areas were calculated from a single visit. Since the coefficient of variation (CV) for this area measurement method was around 6%, eyes with percentage area difference less than the CV were excluded to avoid measurement noise.

### Patient histology

The eye from the CHM donor was fixed in 10% formalin upon enucleation and shipped to the Wilmer Eye Institute. After removing the anterior chamber, a piece was cut nasal to the disc and fixed in 2.5% gluteraldehyde and 2% paraformaldehyde in 1M cacodylate buffer at 4°C for over 24 hours. Tissue was then processed for TEM as previously described [9]. TEM images were collected using a Hitachi H7600 transmission electron microscope at 80KV.

### Statistical analysis

OCT/OCT-A statistical analyses was performed on SPSS v. 25.0 (IBM Corporation) and Microsoft Excel 2017 (Microsoft Corporation). Data are presented as population mean ± standard deviation (SD) or median. Wilcoxon-signed rank test was used to assess whether the difference between CC and EZ area is significant. Correlations were investigated using

Spearman’s rank correlation coefficient (*rho*). All other statistical analyses were performed using GraphPad Prism 8 and data are expressed as mean ± SEM. For comparison between two groups, data were analysed using Students t-test. For grouped analyses, two-way ANOVA with Sidaks multiple comparison test was used. P value of ≤ 0.05 was considered significant.

## Results

### Melanogenesis in *chm ^ru848^* zebrafish

The ultrastructure of the choroid was examined using TEM in wt and *chm^ru848^* zebrafish (Figure 1). Onset of rapid retinal degeneration in *chm^ru848^* fish occurs from 4.5 dpf [7], therefore timepoints of 4 dpf and 5 dpf were selected to analyse the choroidal phenotype. In wt fish, melanosome populations in both RPE and choroidal layers are fully established from 4 dpf. Signs of progressive apoptosis and degradation of melanosomes are observed in the *chm^ru848^* RPE layer, indicated by fewer and smaller melanosomes. A significant absence of melanosomes was observed in the *chm^ru848^*choroidal layer at day 4 and 5. Quantification of total melanin levels in zebrafish eyes revealed melanin was significantly reduced to 70.8 ± 6.4% at 4 dpf (p=0.0105) and 71.3 ± 5.4% at 5 dpf (p=0.0059) in *chm^ru848^* eyes compared to wt (Figure 1C). Expression of genes in the melanogenesis pathway were analysed by RT-qPCR. Expression of *tyr, tryp1a, dct, mitfa, pmela* and *pmelb* were all significantly reduced in the *chm^ru848^*fish from 4 dpf (Figure 1D).

**Figure 1:**
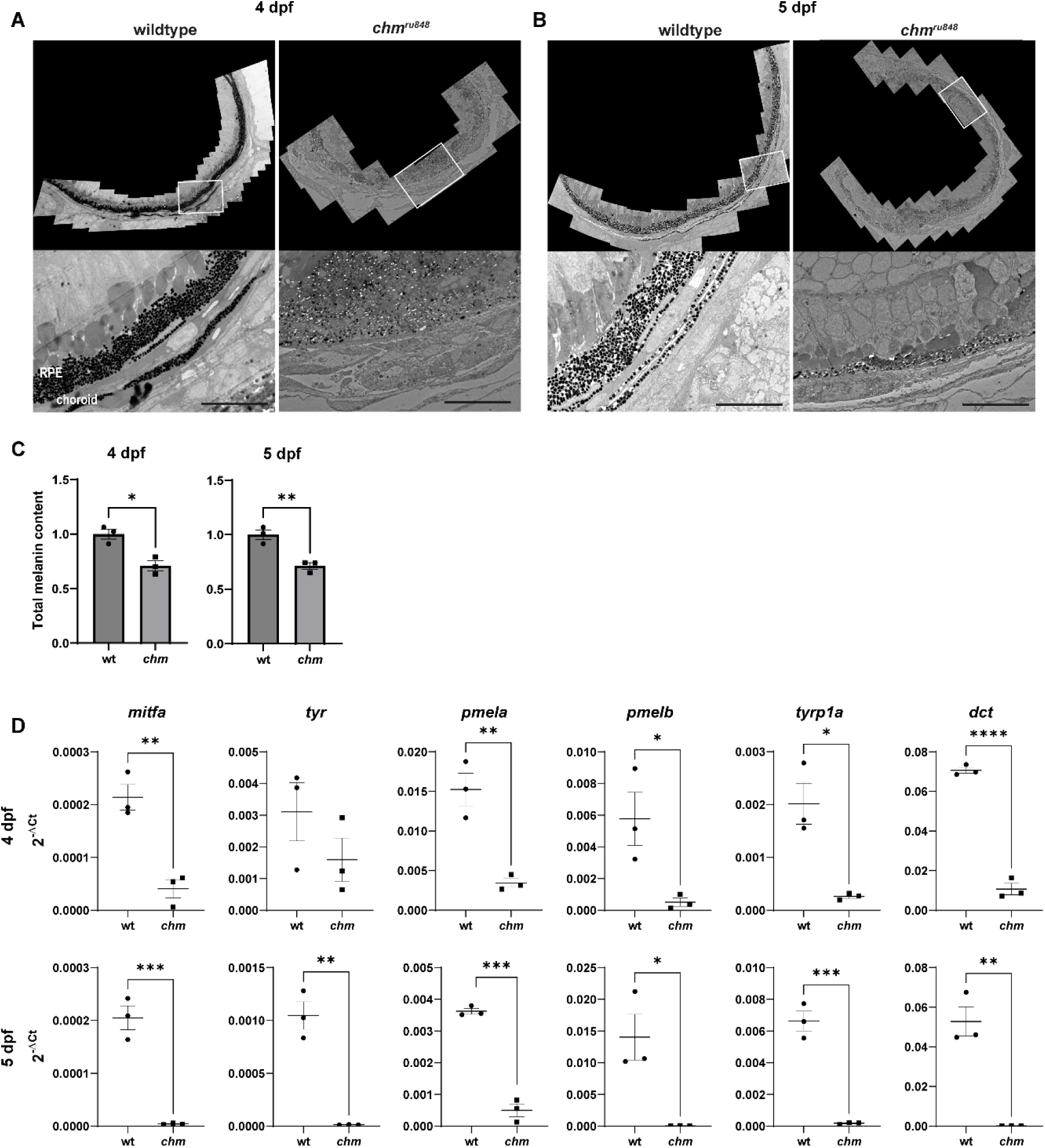
Reduced melanogenesis in *chm ^ru848^*zebrafish. TEM of wt and *chm^ru848^* RPE and choroid at (A) 4 and (B) 5 dpf. Areas in the white boxes have been magnified in the images below. Significantly reduced melanosomes in the RPE layer and a lack of expected choroidal melanosomes was observed in *chm^ru848^*fish from 4 dpf. Scale bar=10µm. (C) Total melanin levels were quantified in zebrafish eyes at 4 dpf and 5 dpf. Melanin levels were significantly reduced in *chm^ru848^* fish at 4 and 5 dpf. (D) Expression of melanogenesis genes were analysed by RT-qPCR at 4 dpf and 5 dpf in zebrafish eyes. Data are expressed as mean ± SEM from n=3. Statistical significance determined by t-test. *p≤0.05, **p≤0.01, ***p≤0.001.

### Retinal microangiography in *chm ^ru848^* zebrafish

The Tg(fli1a:EGFP) line, which expresses EGFP in endothelial cells was crossed with *chm^ru848^* zebrafish and used to analyse choroidal vasculature (Figure 2) using wholemount fluorescent retinal microangiography. In general, CC in *chm^ru848^* fish appeared degenerative, immature and underdeveloped including multiple single endothelial cells not participating in vessel formation at 4 dpf and collapsed and/or disconnected vessels at 5 dpf which did not contribute to forming a functional vasculature, suggesting progressive CC degeneration and loss of function from 4 to 5 dpf. CC vessel density was comparable between wildtype (wt) and *chm^ru848^* fish at 33% and 31% respectively at 5 dpf. Vessel diameter in wt fish at 4 dpf was 8.8±0.4 µm, whereas in *chm^ru848^*fish it was significantly reduced at 6.8±0.5 µm (p=0.0078) (Figure 2C). At 5 dpf the vessel diameter was increased to 10.8±0.7 µm in wt fish, whereas in *chm^ru848^*fish it was significantly lower at 6.6+0.5 µm (p<0.0001). Intussusceptive angiogenesis is the process of new vessel growth via splitting from an existing vessel. Endothelial cells from opposite sides of a capillary wall protrude into the lumen to make contact and the newly formed interstitial pillars (ISP) extend to eventually split the vessel [14]. The number of ISP were counted as a measure of intussusception and found to be significantly reduced in *chm^ru848^* compared to wt fish at 4 dpf (p=0.024) and 5 dpf (p<0.0001) (Figure 2C-D).

**Figure 2:**
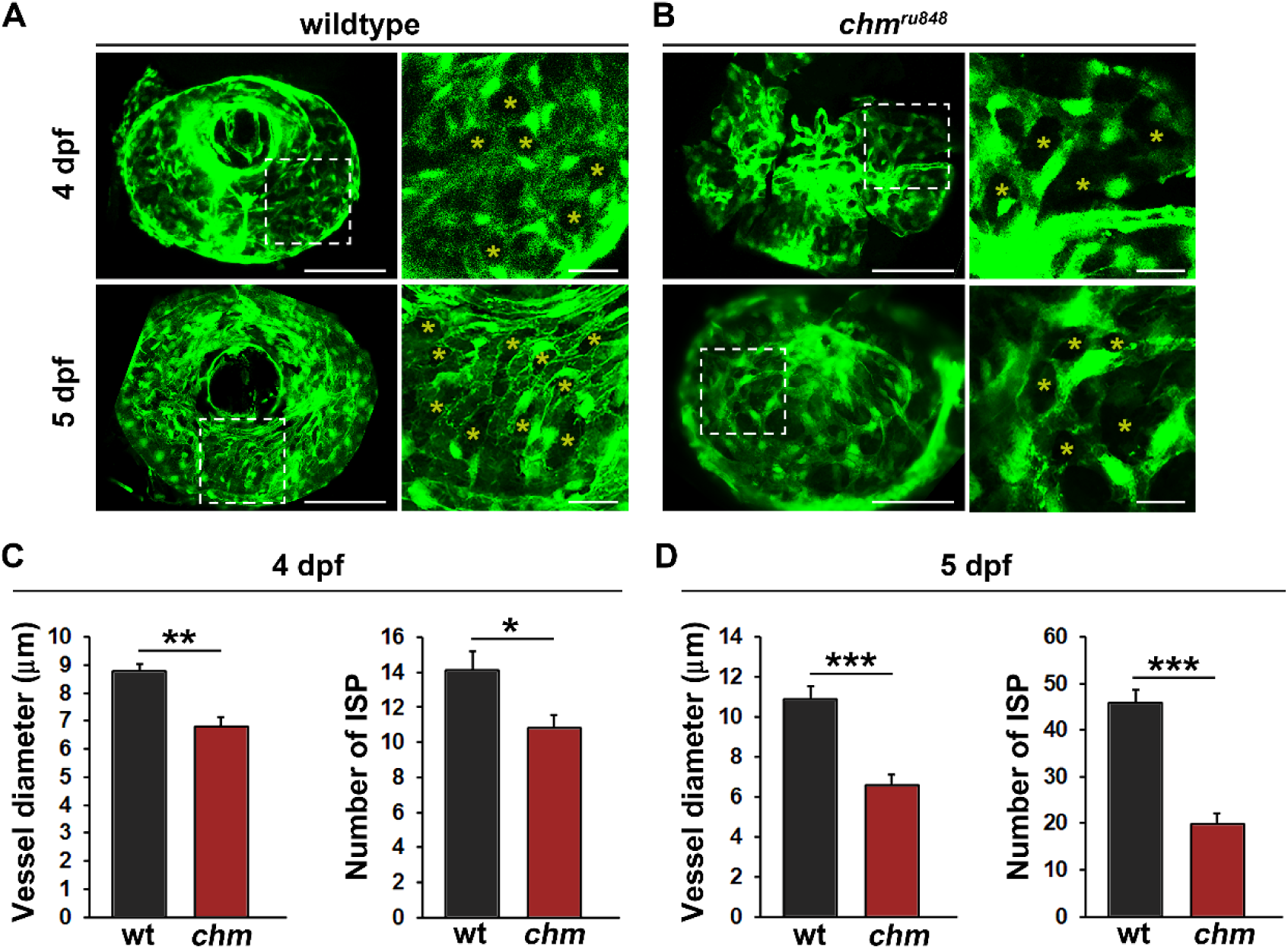
Progressive choriocapillary degeneration in *chm^ru848^* zebrafish larvae. Fluorescent micrographs of the choriocapillaris of fli1a:EGFP;wt (A, wildtype) or fli1a:EGFP;*chm^ru848^* (B, *chm^ru848^*) zebrafish at 4 or 5 dpf. Yellow asterisks indicate interstitial pillars (ISPs). Areas in the white dashed boxes to the left have been magnified in the images to the right. Scale bars indicate 50 µm in the overview images and 10 µm in the magnified images. Quantification of vessel diameter and the number of ISPs at (C) 4 dpf or (D) 5 dpf from the experiment shown in A,B. n=6, 7, 11 and 12 larvae in the wt-4dpf, *chm*-4dpf, wt-5dpf and *chm*-5dpf groups respectively. *:p<0.05, **:p<0.01, ***:p<0.001.

### Choroidal ultrastructure in *Chm^null/WT^* mice

Choroidal histology was examined at 1 month and 1 year in WT (*Chm^flox/flox^*) and *Chm^null/WT^* mice. At 1 month, there was no significant difference in thickness of the choroidal layer between WT and *Chm^null/WT^*mice, however at 1 year, the choroidal layer was significantly thicker in the *Chm^null/WT^* mice (p=0.022) (Figure 3). The ultrastructure of the choroid was then examined using TEM and the area of melanosomes was quantified (Figure 4). Average size of melanosomes in WT choroid at 1 month was 0.110±0.002 µm^2^, but in *Chm^null/WT^* mice was significantly reduced to 0.082±0.001 µm^2^ (p<0.0001). At 1 year, average size of choroidal melanosomes in WT mice was 0.145±0.002 µm^2^, but in *Chm^null/WT^* mice was marginally smaller at 0.141±0.002 µm^2^ (p=0.0123) (Figure 4C).

**Figure 3:**
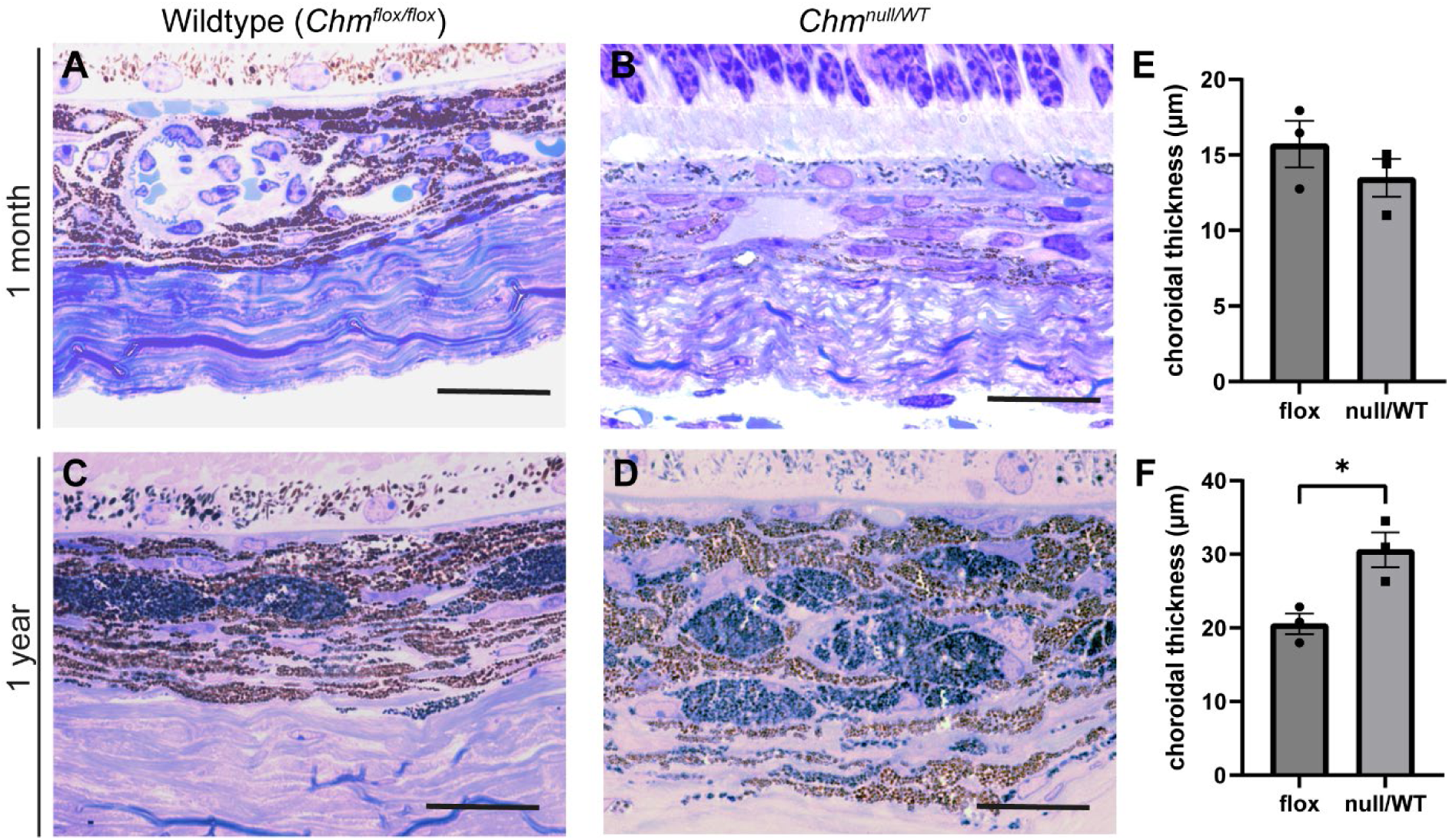
Increased choroidal thickness in *Chm^null/WT^* mice at 1 year. (A-D) Histology of WT(*Chm^flox/flox^*) and *Chm^null/WT^* mouse choroid at 1 month and 1 year. Scale bar=20µm. Thickness of choroid was measured at (E) 1 month and (F) 1 year. Choroid was significantly thicker at 1 year. Data are expressed as mean ± SEM from n=3. Statistical significance determined by t-test. *p≤0.05.

**Figure 4:**
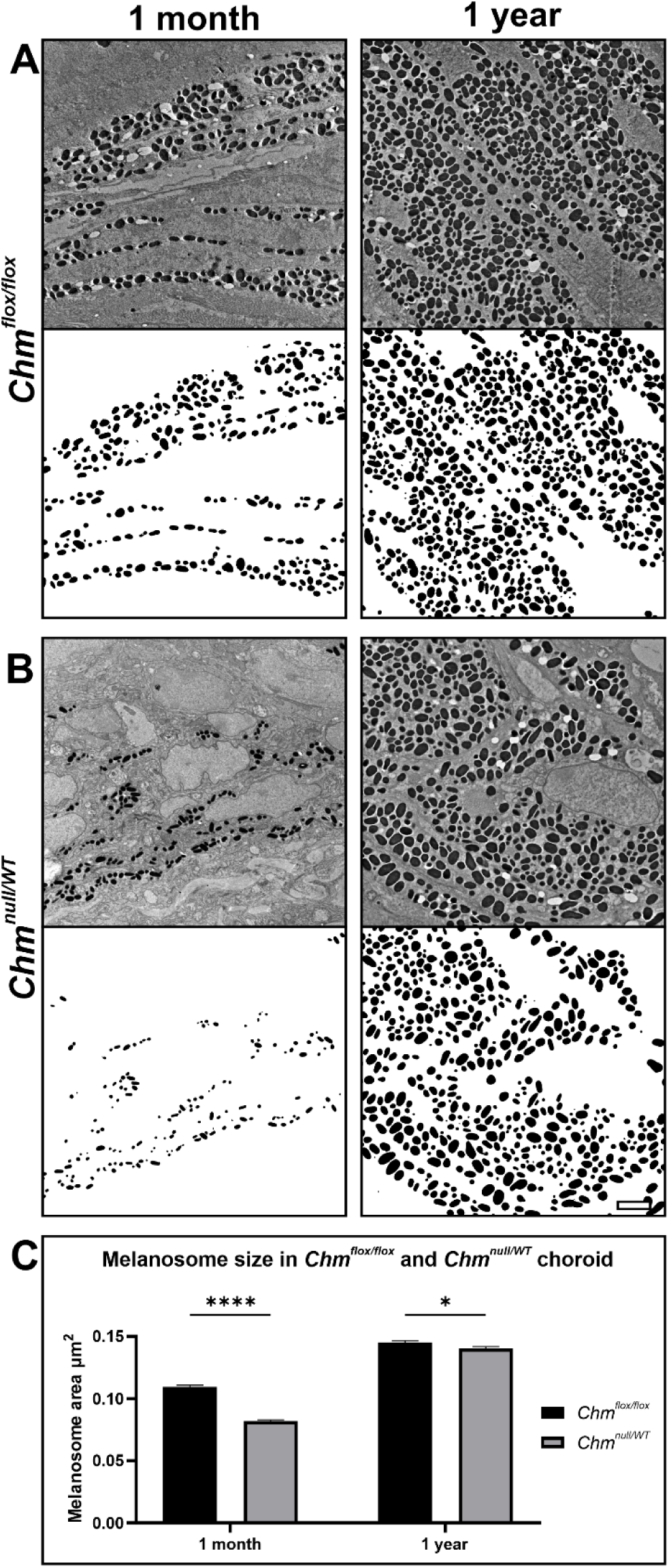
Smaller choroidal melanosomes detected in chm mice at 1 month. (A-B) Electron microscopy of the ultrastructure of WT(*Chm^flox/flox^*) and *Chm^null/WT^* mouse choroid at 1 month and 1 year. Contrast enhanced images of the choroid were used to measure melanosome size using imageJ. (C) Melanosomes were significantly smaller in size in *Chm^null/WT^* mice compared to WT at 1 month. Data are expressed as mean ± SEM. A minimum of 4000 melanosomes were measured from 3 mice per timepoint. Statistical significance determined by two-way ANOVA. *p≤0.05, ****p≤0.0001.

RPE and choroid were dissected and the level of eumelanin and pheomelanin was evaluated by HPLC and mass spectrometry. The characteristic peaks for pheomelanin were not detected in 1 month or 1 year samples, indicating that pheomelanin content is below the level of detection. There was no significant difference in eumelanin content between WT and *Chm^null/WT^* mice at 1 month, however at 1 year eumelanin was reduced to 53±17% in *Chm^null/WT^*RPE/choroid (Figure 5).

**Figure 5:**
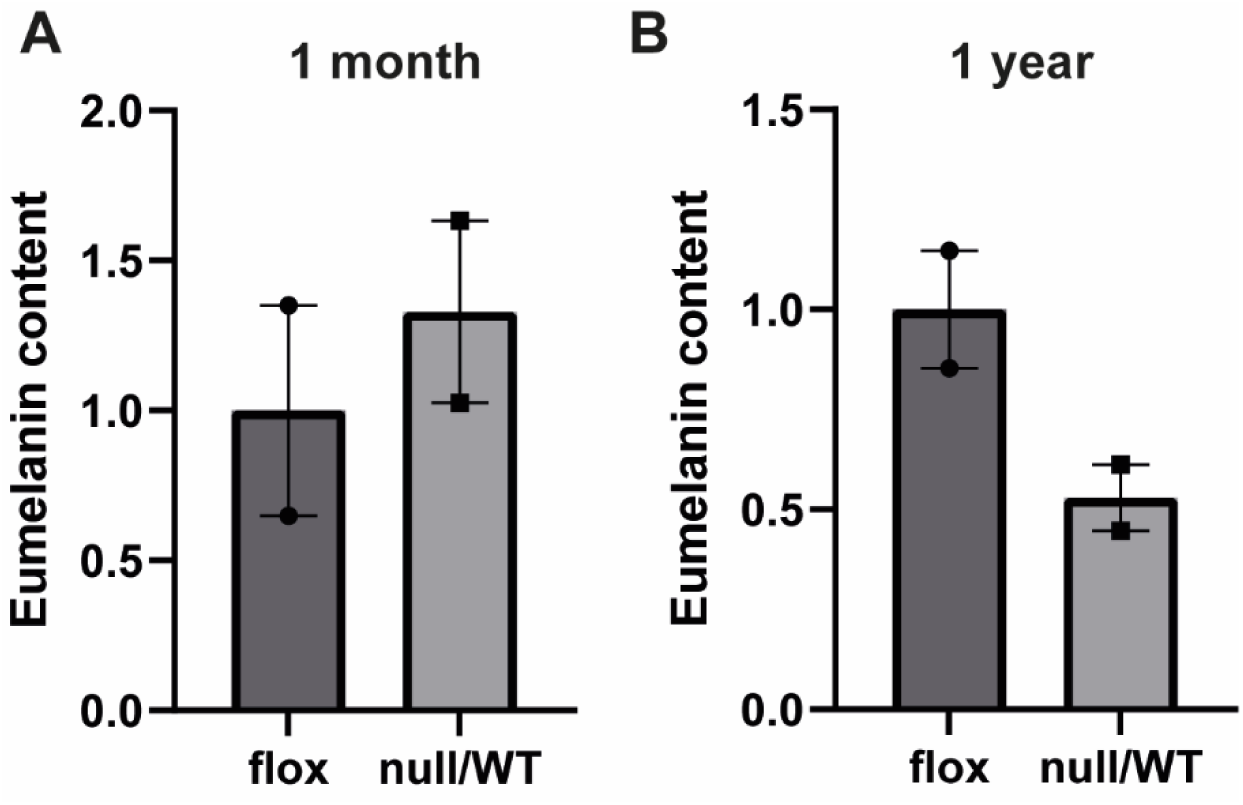
Reduced eumelanin in RPE/choroid of *Chm^null/WT^* mice at 1 year. Levels of eumelanin and pheomelanin were quantified in mice RPE/choroid at (A) 1 month and (B) 1 year by HPLC. Pheomelanin could not be detected in either WT (*Chm^flox/flox^*) or *Chm^null/WT^*mice. Eumelanin levels were reduced in *Chm^null/WT^* mice at 1 year. Data expressed as mean ± SEM from n=2.

### Optical coherence tomography angiography (OCT-A) in CHM patients

Twenty-three eyes from 12 patients (mean age ± SD, 45.28 ± 16.87 years; range, 20-75 years) had gradable OCT/OCT-A images in at least one visit during the natural history study. Eighteen eyes from 11 subjects were included in this analysis. The list of included patients with their genetic results and area measurements is summarised in Table 1. Strong and significant correlation was observed between preserved areas of CC and ellipsoid zone (EZ, the inner/outer segment of photoreceptors) (*rho* = 0.94, *p* < 0.001, Spearman’s rank). However, CC was more severely affected than the EZ in 16 eyes (89%), with mean preserved area difference (± SD) of −0.81 (± 1.31) mm^2^ (median percentage difference, −15.8%, *p* = 0.016, Wilcoxon-Signed rank) (example shown in Figure 6). Although the areas of CC and EZ correlated significantly with patient age (*rho* = −0.55 and −0.49, *p* = 0.034 and 0.04, respectively), no association was observed between age and the absolute or percentage differences between CC and EZ areas.

**Figure 6:**
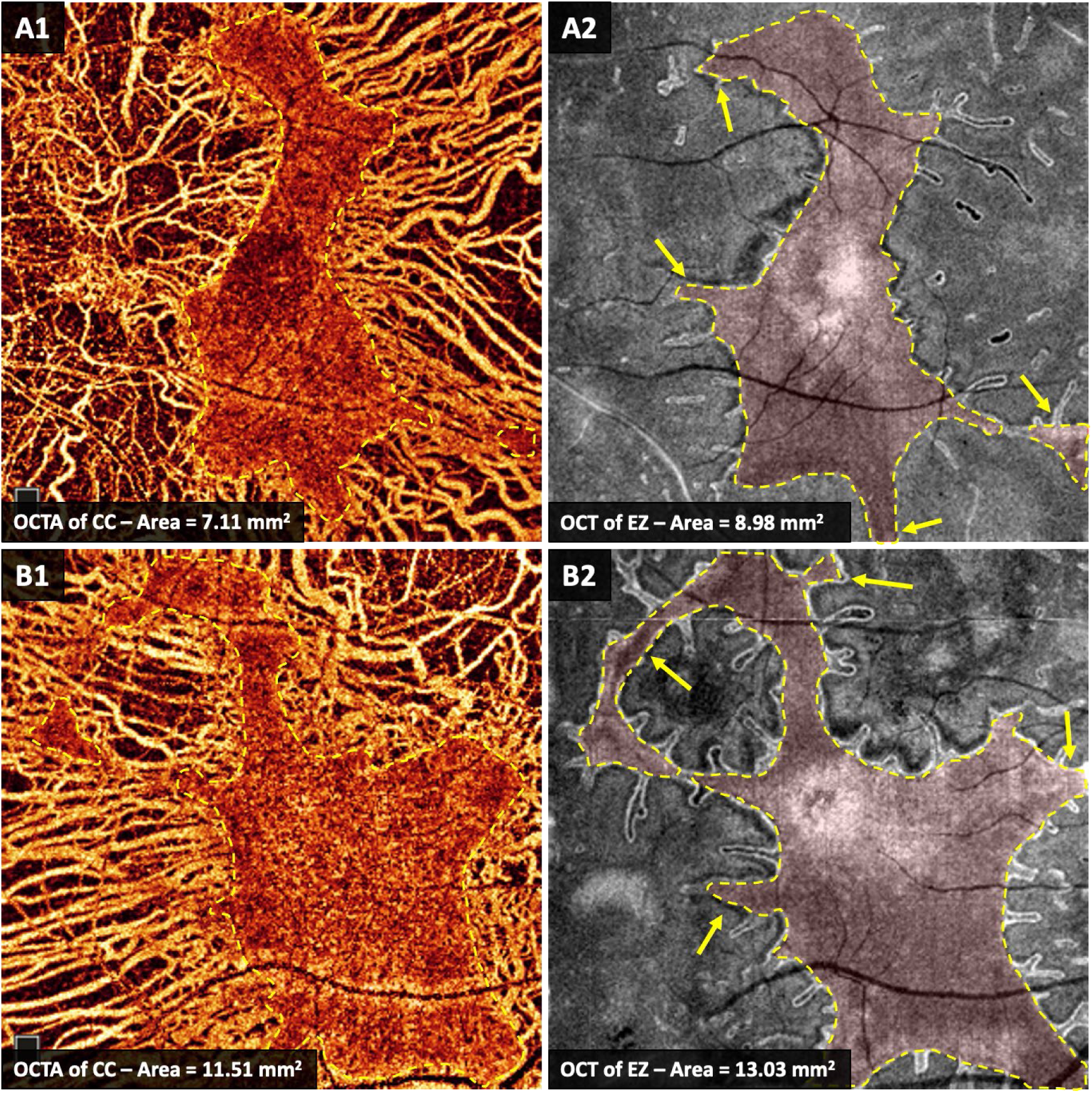
Optical coherence tomography (OCT) and OCT angiography (OCT-A) of the ellipsoid zone (EZ) and choriocapillaris (CC), respectively. (A): The left eye of patient 018. (B): The right eye patient 014. Yellow arrows in A2 and B2 demonstrate areas with preserved EZ where the underlying CC appear degenerated on OCT-A images in A1 and B1.

**Table 1.** Genetic results and area measurements of preserved choriocapillaris (CC) and photoreceptors in choroideremia patients.

| **Pedigree** | **Subject** | **Mutation nucleotide** | **Mutation protein** | **Eye** | **Age** | **CC Area** | **EZ Area** | **Area Difference** |
| --- | --- | --- | --- | --- | --- | --- | --- | --- |
| A | **Patient 001** | c.126C>G | p.Tyr42* | OD | 29 | 1.36 | 1.57 | -0.21 |
|  |  |  |  | OS |  | 1.99 | 2.27 | -0.29 |
| B | **Patient 003** | c.715C>T | p.Arg239* | OS | 62 | 2.95 | 3.17 | -0.21 |
| B | **Patient 015** | c.715C>T | p.Arg239* | OS | 62 | 9.45 | 10.75 | -1.31 |
| C | **Patient 005** | c.1347C>G | p.Tyr449* | OS | 58 | 2.13 | 0.89 | 1.24 |
| D | **Patient 007** | c.698C>G | p.Ser233* | OD | 50 | 0.32 | 0.53 | -0.21 |
|  |  |  |  | OS |  | 0.03 | 0.24 | -0.20 |
| E | **Patient 010** | c.715C>T | p.Arg239* | OS | 42 | 16.83 | 15.62 | 1.21 |
| F | **Patient 011** | c.715C>T | p.Arg239* | OD | 29 | 12.39 | 15.31 | -2.92 |
|  |  |  |  | OS | 28 | 12.95 | 15.01 | -2.06 |
| G | **Patient 013** | c.715C>T | p.Arg239* | OD | 75 | 1.46 | 3.27 | -1.81 |
|  |  |  |  | OS | 74 | 1.55 | 2.62 | -1.07 |
| H | **Patient 014** | c.757C>T | p.Arg253* | OD | 20 | 11.51 | 13.03 | -1.53 |
|  |  |  |  | OS | 21 | 12.87 | 11.14 | 1.73 |
| I | **Patient 017** | c.715C>T | p.Arg239* | OD | 44 | 4.90 | 5.86 | -0.95 |
|  |  |  |  | OS |  | 9.36 | 10.83 | -1.48 |
| J | **Patient 018** | c.799C>T | p.Arg267* | OD | 49 | 11.29 | 13.92 | -2.62 |
|  |  |  |  | OS | 49 | 7.11 | 8.98 | -1.87 |

Areas were measured in mm<sup>2</sup>. CC: choriocapillaris, EZ: ellipsoid zone. Pedigree and subject IDs match those in Hagag et al. (2021).[13]

### Human choroid transmission electron microscopy (TEM)

We expanded our analysis of the ultrastructure of an enucleated eye from a 74-year-old CHM patient [9]. In an 81-year-old age-matched control donor eye, a uniform RPE monolayer is observed beneath photoreceptor segments, and with CC below the Bruch’s membrane. In our initial report of the CHM donor, we observed only areas where either RPE and CC were both lost or preserved. However, closer examination of our TEM blocks revealed isolated areas with RPE still present but no CC (Figure 7). RPE in this area does not form a monolayer and lacks the normal RPE uniform morphology indicative of its disease state.

**Figure 7:**
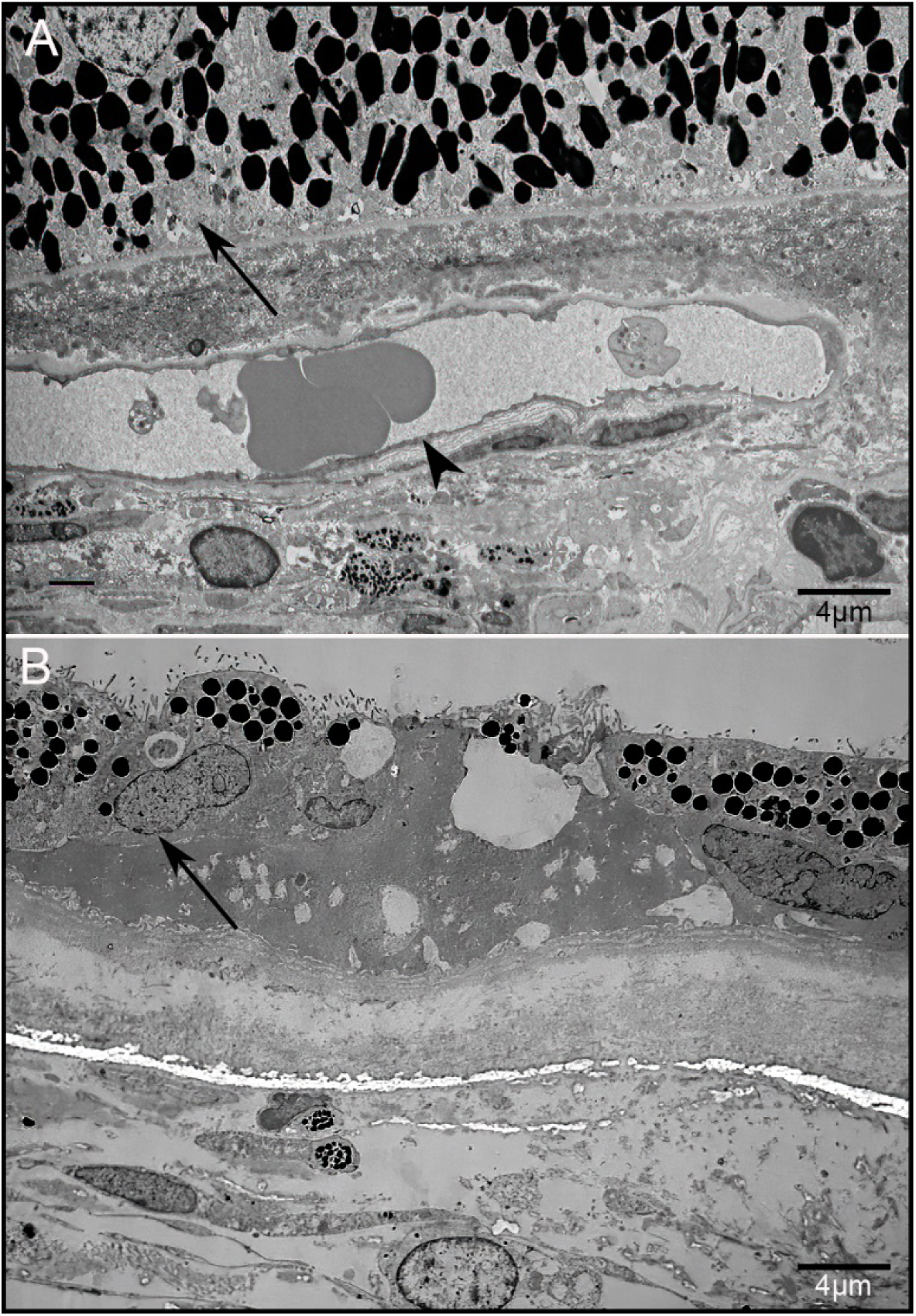
Transmission electron microscopy (TEM) of healthy control and CHM donor eye. (A) TEM micrograph of an 81-year-old healthy control shows an intact RPE monolayer (arrow) with CC vessels below (arrowhead). (B) In the CHM donor eye, only isolated RPE was observed without any underlying CC (arrow). The RPE was not present in a monolayer or displaying uniformity. Scale bar – 4 µm.

## Discussion

The involvement of the choroid in disease progression of CHM is not fully understood; in this study, we examined the ultrastructure of the choroid in zebrafish and mouse models of CHM and found choroidal melanogenesis to be disrupted. Analysis of the choroid in *chm ^ru8484^* zebrafish from 4 dpf revealed severely diminished pigmentation in the choroidal layer, with a complete absence of melanosomes. Overall melanin levels and expression of melanogenesis genes in the *chm ^ru8484^*retina was also significantly reduced from 4 dpf. Smaller melanosomes were detected in the *Chm^null/WT^* mouse choroid at 1 month, with reduced eumelanin levels in the RPE/choroid at 1 year. Rab proteins play a key role in melanogenesis, from melanosome biogenesis to transport [15]. Melanosome development occurs in 4 stages from an unpigmented vesicle to a mature pigmented melanosome [16]. Stage 1 melanosomes do not possess the melanogenic enzymes *TYR*, *TYRP1*, *TYRP2*, therefore melanin synthesis is dependent on trafficking of melanin precursors and enzymes to melanosomes. Rab38 and Rab32 were reported to be involved in trafficking of melanogenic enzymes from the Golgi to melanosomes [17]. Mature melanosomes are transported along the actin cytoskeleton, facilitated by Rab27a [1]. Lack of REP1 results in underprenylation of Rab proteins; as several Rab proteins are involved in melanosome biogenesis, loss of REP1 can be expected to cause disruption to melanogenesis. Using the *Cht* mouse model, which lacks functional Rab38, Lopes et al. showed that the majority of melanosomes are synthesised before birth in the RPE, however, in the choroid, most melanosomes are synthesised after birth, with a low level of melanosome biogenesis occurring in the adult choroid [18]. Patchy depigmentation of the RPE was reported from embryonic stage P7 in *Chm^null/WT^* mice [8], and reduction in melanin synthesis genes was observed in the *chm* zebrafish from 4 dpf, indicating REP1 may play an important role during development. Melanosomes of the RPE are derived from the neuroectoderm, whereas choroidal melanosomes are derived from the neural crest, therefore different Rab proteins may be required for melanosome biogenesis in each layer and may be differentially prenylated by REP1. For example, in mice lacking functional Rab38, the RPE is severely depigmented however pigmentation in the choroid is only mildly affected [18]. Further study is required to fully understand the role of REP1 in melanogenesis.

In addition to melanogenesis, vasculogenesis was also disrupted in *chm ^ru8484^* zebrafish. In *Mitf^mi-bw^* mice lacking choroidal melanocytes, but with a pigmented RPE, the choroid was thinner and choroidal vasculature was disrupted, suggesting that choroidal melanocytes are important for maintaining the normal vasculature of the choroid [19]. Melanocytes were also found to secrete fibromodulin, a potent angiogenic factor, which stimulates blood vessel formation [20]. Reduced number of ISP were detected in *chm^ru8484^* fish, indicating impaired melanogenesis leads to disrupted angiogenesis with decreased ability to grow new vessels.

CHM patients display a thinner choroid in the natural history of the disease [13, 21], however we observed a significantly thicker choroid in *Chm^null/WT^* mice at 1 year. A thicker choroid can be associated with inflammation. Infiltration of glial cells was reported in the retina of 6-month-old *Chm^null/WT^* mice [22]. In histopathological examination of a 30 year old CHM patient, mild inflammatory cellular infiltration was detected in the choroid, along with inflammatory T-lymphocytes surrounding choroidal vessels [23]. Glial cell migration through breaks in the Bruchs membrane was reported in a 66 year old CHM patient [24] and inflammatory cells were also detected in the choroid of a 91-year-old female carrier [25]. Recently, human choroidal melanocytes were shown to respond to inflammation by increased expression of inflammatory cytokines and genes associated with cell adhesion and angiogenesis [26]. Impaired melanogenesis may impact ability of the choroid to respond to inflammation, leading to increased inflammation and a thicker choroid.

At 1 year, although the choroid was thicker, eumelanin levels were reduced in the *Chm^null/WT^* RPE/choroid, which corresponds with patchy depigmentation of the RPE, that has been reported in these mice from 9 months [8]. Eumelanin, which is known to be a potent antioxidant and free radical scavenger, was also reduced in zebrafish from 4 dpf. We recently reported increased levels of oxidative stress related metabolites in CHM patient plasma [27]. Reduced melanin levels may contribute to a decreased ability to protect against reactive oxygen species, leading to increased oxidative damage in the retina, highlighting the use of antioxidants as a possible therapeutic avenue.

Most studies have suggested that the initial cause of degeneration in CHM is due to early involvement of the RPE [21, 28–32], with little focus on the choroid. We therefore examined the choroid in CHM patients using OCT/A and found the area of preserved CC was smaller than that of the EZ, indicating that the choroid is degenerating at a faster rate than the photoreceptors. In a study by Arrigo et al., reduced vessel density of the deep capillary plexus and CC was reported in CHM patients [33] and Murro et al. reported reduced vascular density of the CC in young patients with preserved FAF and preserved hyper-reflective outer retinal layers [34]. Histopathologic examination of donated eyes from a 66-year-old male CHM patient with extensive chorioretinal atrophy revealed hypoproduction of basement membranes by the vascular endothelial cells of the choroidal and iris vessels, suggesting a primary defect within the vessel walls itself, leading to secondary loss of the adjacent RPE [24]. We recently reported loss of CC associated with RPE degeneration and migration into the retina in all but the far periphery in the eyes of two CHM donors. In the far periphery, both RPE and CC remained [9]. In the present report, we revisited the TEM blocks of one CHM donor and found the presence of abnormal RPE anterior to missing CC indicating that CC loss may occur independently of RPE damage. In the very least, this data demonstrates that CC loss occurs in close succession to RPE loss or damage. It is important to consider, however, that any changes to RPE cells could result in increased cytokine production as well as altered VEGF production and altered metabolic activity. These could all impact CC health. At the same time, we cannot rule out the possibility that the RPE cell changes result from a loss of CC in this area, an idea supported by clinical and animal model data presented herein.

A number of clinical trials for AAV gene therapy via subretinal injection have taken place, however recent results from the phase III clinical trial reported failure to meet primary and secondary endpoints (https://investors.biogen.com/news-releases/news-release-details/biogen-announces-topline-results-phase-3-gene-therapy-study). Currently, a phase I/II clinical trial for intravitreal injection of recombinant adeno-associated virus (AAV) gene therapy, 4D-110, is underway (NCT04483440). However, future therapeutics should consider also targeting the choroid in order to counter its degenerative processes, which may be influencing the RPE changes. This may be through systemic administration (oral or intravenous) or through suprachoroidal delivery.

In summary, pigmentary disruptions in CHM animal models reveal an important role for REP1 in melanogenesis, with further impact on vasculogenesis and modulation of inflammatory response. Further work is required to fully understand the development of the choroid in patients with CHM and the complex relationship between REP1, Rabs and melanogenesis. However, potential therapies that improve melanin production may be a new therapeutic target to restore function and protect the retina, hence slowing the impact of chorioretinal degeneration in CHM.

## Acknowledgements

This research was funded by Wellcome Trust, grant number 205174/Z/16/Z, the Choroideremia Research Foundation, USA and Moorfields Eye Charity.

## Supplementary material

### Zebrafish qRT-PCR primers

**Table S1:**
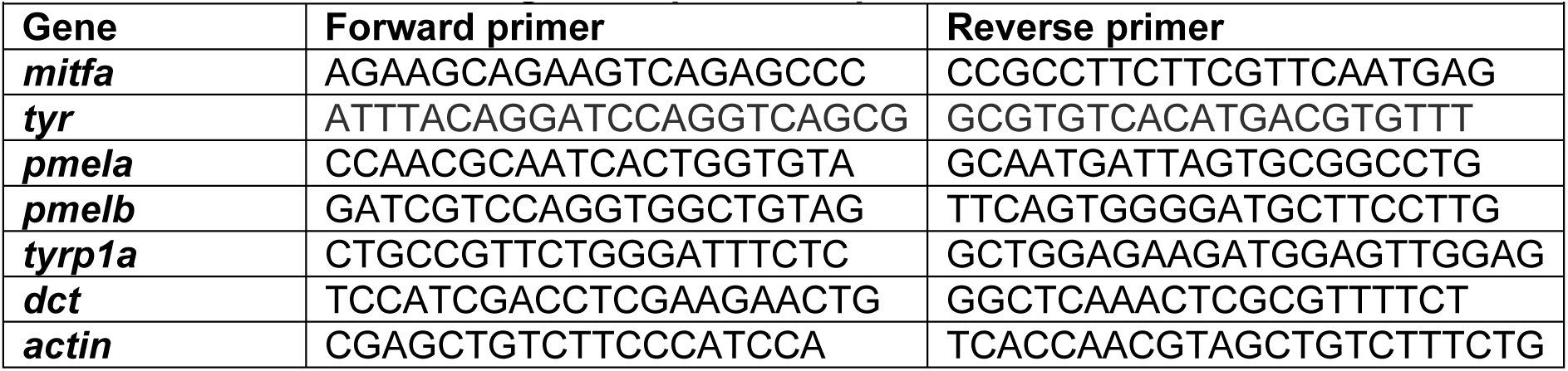
Zebrafish melanogenesis primer sequences.

